# Dichotomous transcriptome divergence at the cell-type level between the *Drosophila* male and female germlines

**DOI:** 10.1101/2025.07.04.663238

**Authors:** Imtiyaz E. Hariyani, Sudeshna Das, Emma M. Le, Carmen Gamero-Castano, Tina Soroudi, Joshua Choi, Vivek Swarup, Jose M. Ranz

## Abstract

Gene expression evolution in reproductive organs plays a central role in species divergence, yet cell-type-resolved patterns in invertebrates remain poorly understood. We used single-nucleus RNA-sequencing to profile testis and ovary transcriptomes from the sibling species *Drosophila melanogaster* and *D. simulans*. Despite conserved cellular composition, we observed highly cell-type-specific expression and coexpression network divergence between species. In the testis, mitotic cells were conserved across species, mirroring patterns in mammals, while divergence peaked in late spermatocytes. In the ovary, divergence was less pronounced, peaking in early germline and late-stage follicle cells, and enriched on the *X* chromosome, consistent with a faster-*X* effect driven by positive selection. Genes expressed in these cell types exhibited narrower expression breadth, younger phylogenetic age, and elevated rates of protein evolution in both tissues. Our findings reveal contrasting evolutionary regimes in male and female germlines, shaped by adaptive and non-adaptive mechanisms, contingent on cell type and chromosomal context.

---

Gene expression changes are a key driver of species adaptation and phenotypic diversification ^1–4^. While early comparative transcriptomic studies using microarrays or bulk RNA-sequencing helped characterize large-scale expression differences at the tissue level, they lacked cell type resolution and were confounded by differences in tissue composition between species ^5,6^. Single-cell and single-nucleus RNA sequencing (scRNA-seq and snRNA-seq, respectively) overcome these limitations by enabling direct comparisons of homologous cell types at unprecedented granularity within complex tissues ^7,8^. These approaches have led to more precise evolutionary inferences—for example, in mammalian tissues such as the testis ^5,9^ and various regions of the brain ^10,11^—by clarifying which cell types are most conserved versus those contributing disproportionately to expression divergence.

The gonads are central to understanding expression evolution, as they harbor most sex-biased genes ^12^ and exhibit distinct patterns of gene expression ^13–15^. *Drosophila melanogaster* has served as a premier model organism for studying gametogenesis, primarily using bulk transcriptomic data, often derived from whole-body samples ^16–21^. More recently, single-cell studies in *Drosophila* have revealed stage- and cell type-specific expression patterns across both testis and ovary ^7,22–30^. For example, DNA damage response genes are enriched in early germline stages of both gonadal tissues ^22,27^, and late spermatogenesis is marked by an enrichment of de novo genes ^22^. However, reliance on single-strain datasets has limited our ability to capture the intraspecific genetic diversity of *D. melanogaster* ^31–33^, which is crucial for understanding functional constraints and identifying expression changes that truly contribute to species divergence ^34,35^. Key questions remain unresolved: Which gonadal cell types drive interspecific divergence? Do the testis and ovary follow parallel evolutionary trajectories, or do their constraints and modes of evolution fundamentally differ? To what extent is observed divergence driven by adaptive versus neutral forces? And what functional pathways underlie this divergence?

Despite evolutionary divergence, gametogenesis in *Drosophila* and mammals shares conserved developmental features, including similar differentiation stages and core molecular pathways ^36–38^. In mammals, transcriptomic divergence is not uniform across the testis: it peaks in meiotic and post-meiotic germ cells of the testis, while early germline cells remain highly conserved—likely due to strong functional constraints tied to their pleiotropic roles ^9,39,40^. In the mammalian ovary, the theca cells—endocrine somatic cells that surround the follicle—accumulate greater expression divergence between species compared to other ovarian somatic cell types ^41^. Whether *Drosophila* exhibits similar patterns of stage- or cell-type-specific divergence in either gonadal tissue remains unknown. Comparative single-cell analyses of closely related *Drosophila* species remain scarce, making it difficult to discern whether previously reported expression divergence reflects true biological differences or methodological artifacts such as confounding or compensatory effects ^13,16–18^. If substantial interspecific expression differences exist in *Drosophila*, it is unclear whether they are broadly distributed across cell types or concentrated in terminally differentiated stages, similar to patterns observed in the mammalian testis.

To address these questions, we used snRNA-seq to compare the testis and ovary transcriptomes of three strains spanning two closely related species, *D. melanogaster* and *D. simulans*, which diverged ∼1.4 mya ^42^. We characterized expression divergence across cell types by integrating two complementary analyses: differential expression at the level of individual genes and network divergence at a systems level. We also examined how differentiation trajectories in spermatogenesis and oogenesis relate to pleiotropic constraint, using proxies such as expression breadth and phylogenetic gene age. Our results reveal a dichotomy in the tempo and mode of transcriptome evolution between the male and female germline. Testis transcriptomes evolve rapidly and in a highly cell-type-specific manner, with divergence in meiotic cells driven by a combination of adaptive and non-adaptive forces. In contrast, the ovary exhibits more conserved expression, consistent with stronger pleiotropic constraint—except in a subset of germline and late-stage follicle cells, which show pronounced divergence. We further show that the cell types accumulating divergent genes also tend to have narrow expression breadth and younger phylogenetic age, especially in the testis, and that divergent genes are more likely to be on the *X* chromosome in the ovary. Together, these findings uncover how distinct evolutionary forces shape gene expression programs across cell types in the male and female germlines.

## RESULTS

### Combinatorial barcoding enables cell-type-resolved gonadal transcriptomes across Drosophila *species*

To characterize gene expression divergence between closely related *Drosophila* species at cell-type resolution, we performed single-nucleus RNA-sequencing (Parse Biosciences Evercode v2 ^43^) on gonadal tissues from two strains of *Drosophila melanogaster* (A4, ISO1) and one strain of *D. simulans* (w^501^), selected to capture intra- and interspecific divergence within the *D. melanogaster* species subgroup. We profiled testes from 1–2-day-old naïve males and ovaries from 3–day-old virgin females using split-pool combinatorial barcoding ^43^. After quality control and doublet removal, we retained 106,328 high-quality nuclei–40,274 nuclei from testis and 66,054 nuclei from ovary–across 12 biological samples (3 strains × 2 tissues × 2 biological replicates). To minimize mapping artifacts due to nucleotide differences, we aligned reads to strain-specific genome assemblies. We then integrated cross-strain data for each tissue using Harmony ^44^, which best preserved cell-type and species structure among the tools tested (Methods). With an average of 44,742 reads per nucleus (Supplementary Table 1), our dataset is comparable to or exceeds those of previous studies (Supplementary Tables 2-3), enabling high-resolution comparisons of conserved and divergent expression patterns within and between species.

### Cellular composition of *Drosophila* gonads is conserved across species

We identified conserved germline and somatic cell populations in both testis and ovary, whose proportions and marker expression were highly conserved across all three strains. Using dimensionality reduction (UMAP) and graph-based clustering, we found 10 testis and 17 ovary cell types and annotated them using known marker genes (Supplementary Table 4). In the testis, we identified cells undergoing mitosis (Mi; early and late spermatogonia) and meiosis (Me; early, mid, late, maturing primary spermatocytes, and post-meiotic spermatids), along with three somatic populations (S; epithelial, cyst, and hub cells). In the ovary, we distinguished early germline cells in the germarium (G), somatic cells in the germarium (GS; follicle stem cells, early follicle cells, stalk, and polar cells), and multiple somatic populations in the epithelium (EpS; main body follicle cells, oviduct, ovarian sheath muscle, stretch cells and terminal corpus luteum cells). UMAP visualization faithfully recapitulated the expected spatial arrangement of these cell types, despite the absence of spatial data (Fig. 1a,b). We validated these annotations using developmental trajectory reconstruction, which recovered the expected progression of germline differentiation (Supplementary Fig. 1).

**Figure 1.**
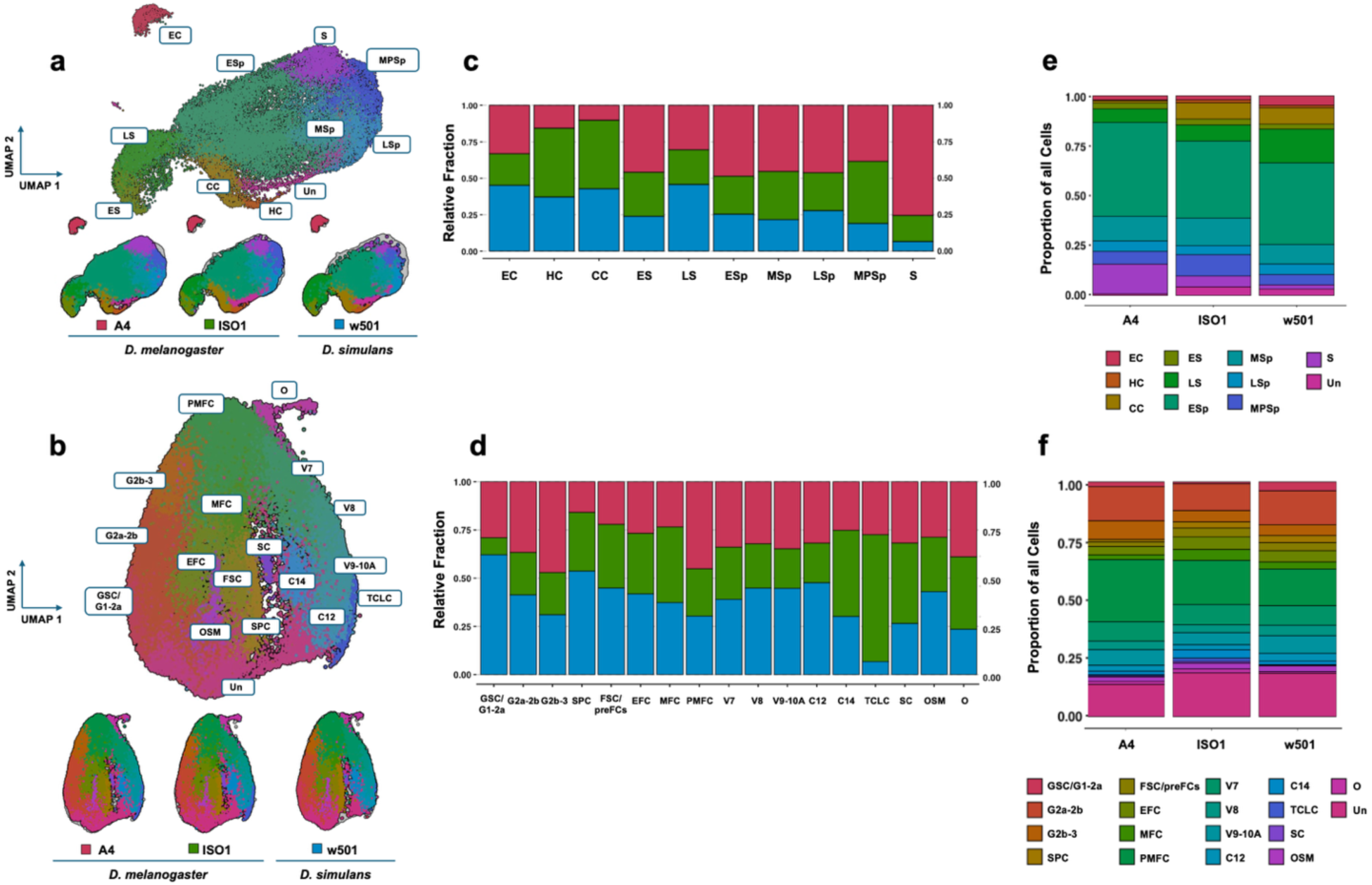
Single-nucleus transcriptomic landscapes of *Drosophila* testis and ovary across strains. **a, b** UMAP visualization of transcriptomic profiles from the testis and ovary, respectively, colored by annotated cell types. Each cluster represents a distinct cell type that corresponds to a particular stage during spermatogenesis and oogenesis respectively, combined for all three strains (above) and individually in *D. melanogaster* strains A4 and ISO1, and *D. simulans* strain w^501^ (below). **c, d** Stacked bar plot showing the relative fraction of identified cell types for each strain in the testis and ovary, respectively. **e, f** Stacked bar plot showing the proportion of all cells annotated for each identified cell type across the three strains in the testis and ovary, respectively. All annotated cell types were included after quality control filtering, regardless of number of nuclei. Testis cell types: EC, epithelial cells; HC, hub cells; CC, cyst cells; ES, germline stem cells and early spermatogonia; LS, late spermatogonia; ESp, early spermatocytes; MSp, mid spermatocytes; LSp, late spermatocytes; MPSp, maturing primary spermatocytes; and S, spermatids. Ovary cell types: GSC/G1-2a, germline stem cells and germarium region 1 and 2a cells; G2a-2b, germarium region 2a and 2b cells; G2b-3, germarium region 2b and 3 cells; SPC, stalk and polar cells; FSC/preFCs, follicle stem cells and pre-follicle cells; EFC, early follicle cells; MFC, mitotic follicle cells stage 1-5; PMFC, post-mitotic follicle cells stage 6; V7, vitellogenic main-body follicle cells (MBFCs) stage 7; V8, vitellogenic MBFCs stage 8; V9-10A, vitellogenic MBFCs stage 9-10A; C12, choriogenic MBFCs stage 12; C14, choriogenic MBFCs stage 14; TCLC, terminal corpus luteum cells; SC, stretch cells; OSM, ovarian sheath muscle; and O, oviduct. Un, unannotated.

The testis and ovary cell types displayed strong signatures of evolutionary stability, with conserved relative proportions across strains (Fig. 1c,d; Supplementary Fig. 2; Supplementary Table 5; Supplementary Note 1) and minimal differences in cell type composition (Fig. 1e,f). Cell type proportions were highly correlated between strains and more similar between the two *D. melanogaster* strains than between either and *D. simulans*, accurately reflecting phylogenetic distance (Supplementary Table 6). Marker gene expression remained remarkably consistent across strains for all annotated cell types (Supplementary Figs. 3 and 4). Principal component analysis revealed that expression profiles clustered by species and by broad cell type category— mitotic, meiotic, and somatic in the testis, and along a developmental trajectory continuum in the ovary (Fig. 2a,b). Biological replicates consistently clustered together, and the two strains of *D. melanogaster* always grouped more closely with each other than with *D. simulans*, reflecting their phylogenetic relatedness and confirming the integrity of our sample preparation and sequencing pipeline. Together, these findings indicate the conserved developmental architecture of gametogenesis, enabling cell-type-level comparisons of gene expression evolution across species.

**Figure 2.**
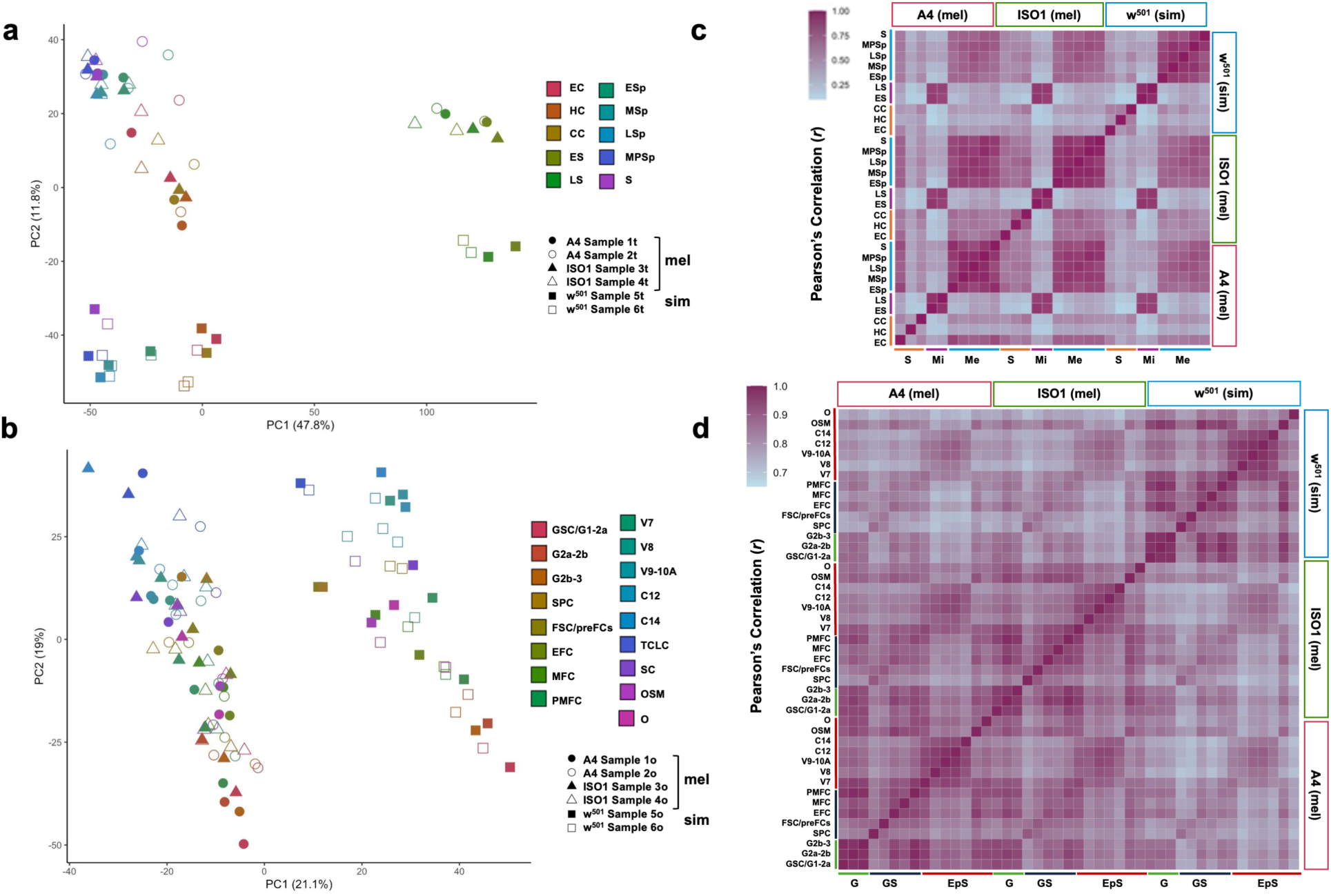
Intra- and inter-specific expression variation and correlations across cell types in *Drosophila* gonads. Principal component analysis (PCA) of cell-type pseudo-bulks. **a** Testis. PC1 separates the mitotic cells from the meiotic and somatic cells. PC2 shows separation between species. **b** Ovary. PC1 shows the separation between species. PC2 shows cell progression along the oogenesis trajectory. Correlation matrices depicting Pearson’s correlation coefficients between the expression levels of genes found expressed in common between any two cell types across the three strains assayed in **c**, testis and **d**, ovary. A4 and ISO1, *D. melanogaster* strains; w^501^, *D. simulans* strain. Broad categories of cell types are indicated: somatic (S), mitotic (Mi), and meiotic (Me) categories for the testis; and germline (G), germarium somatic (GS), and epithelium somatic (EpS) for the ovary. In the correlation analysis, only cell types with at least 50 cells per strain and 350 cells across strains were included; as a result, the TCLC and SC ovarian cell types were excluded. Genes were required to be expressed in at least 1% of cells with a minimum average expression of 0.01. Testis cell types: EC, epithelial cells; HC, hub cells; CC, cyst cells; ES, germline stem cells and early spermatogonia; LS, late spermatogonia; ESp, early spermatocytes; MSp, mid spermatocytes; LSp, late spermatocytes; MPSp, maturing primary spermatocytes; and S, spermatids. Ovary cell types: GSC/G1-2a, germline stem cells and germarium region 1 and 2a cells; G2a-2b, germarium region 2a and 2b cells; G2b-3, germarium region 2b and 3 cells; SPC, stalk and polar cells; FSC/preFCs, follicle stem cells and pre-follicle cells; EFC, early follicle cells; MFC, mitotic follicle cells stage 1-5; PMFC, post-mitotic follicle cells stage 6; V7, vitellogenic main-body follicle cells (MBFCs) stage 7; V8, vitellogenic MBFCs stage 8; V9-10A, vitellogenic MBFCs stage 9-10A; C12, choriogenic MBFCs stage 12; C14, choriogenic MBFCs stage 14; TCLC, terminal corpus luteum cells; SC, stretch cells; OSM, ovarian sheath muscle; and O, oviduct.

### Contrasting patterns of cell type-specific expression divergence in the testis and ovary

The conserved cellular architecture of the gonads extended to the number of genes expressed per cell type, which remained consistent across strains, although we noted sharp within-tissue variation. In the testis, late and maturing primary spermatocytes (LSp and MPSp, respectively), exhibited the highest numbers of expressed genes, whereas mitotic cell types had the fewest (Kruskal-Wallis chi-squared, *P<*0.001 for each strain; Supplementary Table 7 for post-hoc tests). In the ovary, germline stem cells within the germarium 1 and 2a (GSC/G1-2a), and stage 12 choriogenic MBFCs (C12) expressed the highest numbers of genes (Kruskal-Wallis chi-squared, *P<*0.001 for each strain; Supplementary Table 7 for post-hoc tests). These patterns remained stable across all three strains (Supplementary Fig. 5).

Testis and ovary cell types displayed distinct trajectories of transcriptome divergence, with stronger interspecific differences in the testis. To quantify these patterns within (A4 vs. ISO1) and between species (*D. melanogaster* vs. *D. simulans*), we extracted the average expression of each gene for each strain per gonadal cell type (Supplementary Data 1,2) and constructed gene expression correlation matrices for each cell type and tissue (Fig. 2c,d). Intrastrain expression correlations were generally higher between cell types in the ovary than in the testis, indicating that transcriptome divergence in the testis follows a more cell-type-specific trajectory. In both tissues, we observed higher expression similarity among cell types belonging to the same broad cell type category (S, Mi, and Me for the testis; G, GS, and EpS for the ovary). Between strains, expression correlations were higher within species than between species for all categories. In the testis, meiotic populations exhibited the greatest interspecific divergence, whereas mitotic cell types–germline stem cells / early spermatogonia and late spermatogonia–displayed the lowest divergence both within and between species, suggesting a more conserved early transcriptional program. In contrast, the ovary showed higher intra- and inter-specific correlations across most cell types, indicating generally conserved expression patterns across both gametic and somatic categories, except for the late-stage main body follicle cells. Together, these findings reveal contrasting patterns of cell type-specific expression divergence in the gonads, with meiotic types evolving more rapidly in the testis and epithelium somatic cells diverging more in the ovary.

### Differentiating germline cells harbor most intra- and interspecific expression differences

To identify the specific genes driving the expression correlation patterns, we performed differential expression analyses within and between species (Supplementary Data 3,4). We defined differentially expressed genes (DEGs) using a 1% FDR and a log_2_ fold-change ≥ |1|, based on pairwise comparisons at both the intraspecific (ISO1 vs A4) and interspecific (ISO1 vs w^501^, and A4 vs w^501^) levels for each cell type and tissue. To ensure consistent comparisons across strains and species, we restricted our analysis to 11,481 one-to-one orthologous genes ^45^. Consistent with previous work on the interspecific evolution of sex-biased expression ^16,17,20,21^, the testis showed greater expression divergence than the ovary (3,253 DEGs or 35.3% vs 1,748 DEGs or 20.5% of the expressed genes in each tissue, respectively; 2-sample test for equality of proportions with continuity correction or 2STEP, χ^2^=479.2, d.f.=1, *P<*2.2×10^-16^) (Fig. 3a,e).

**Figure 3.**
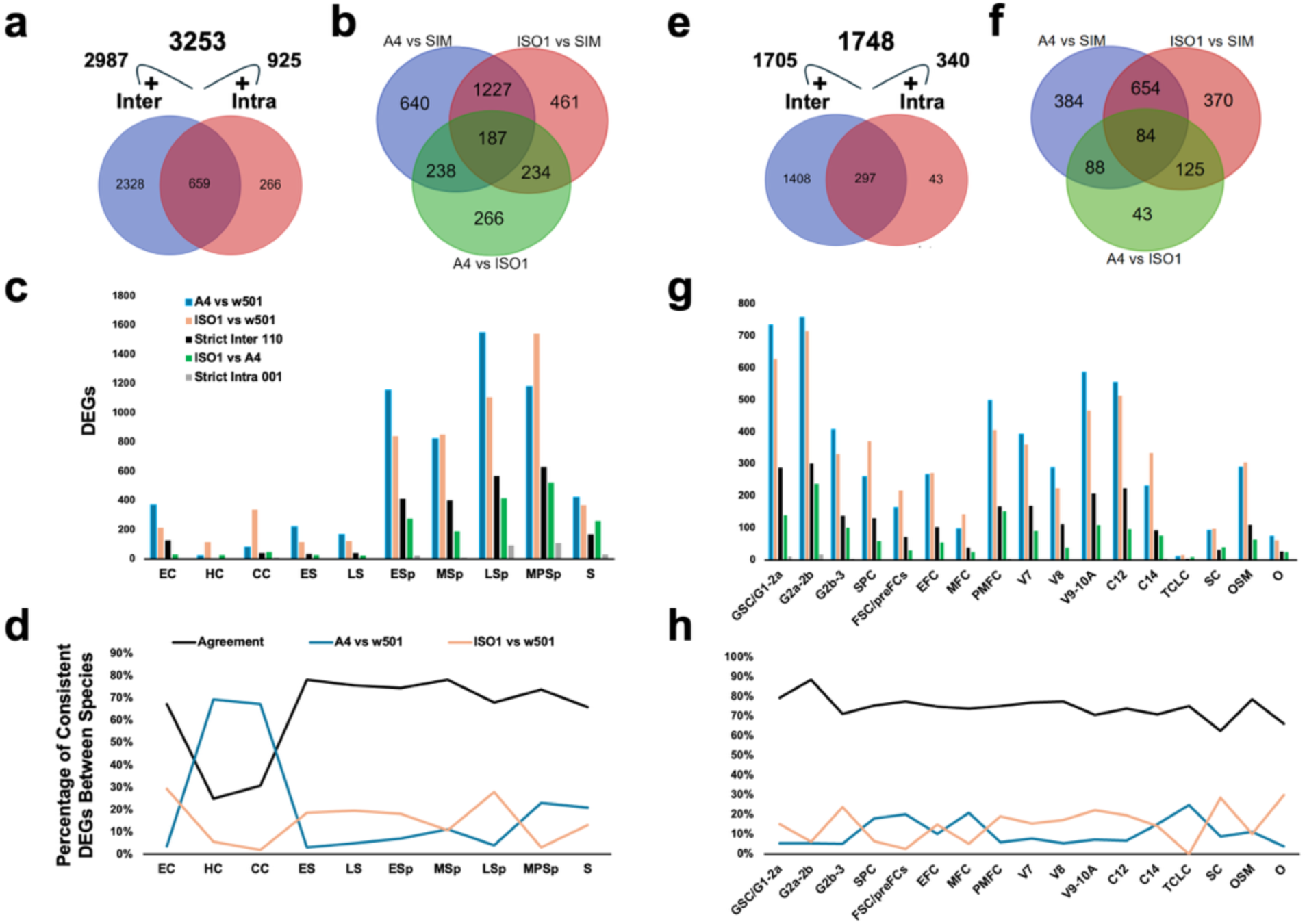
Magnitude of expression differentiation across testis and ovary cell types at two evolutionary timescales. Venn diagrams showing the number of differentially expressed genes (DEGs) unique to, and shared between *D. melanogaster* and *D. simulans* in the testis: **a**, at the inter- and intraspecific levels; and **b**, across the three pairwise comparisons involving the *D. melanogaster* strains A4 and ISO1, and the *D. simulans* strain w^501^. **c**, Patterns of differential expression across pairwise comparisons and testis cell types. Strict Inter (pattern 110): DEGs in both interspecific contrasts in the same cell type without being differentially expressed at the intraspecific level. Strict Intra (pattern 001): DEGs at the intraspecific level without being differentially expressed at the interspecific level. **d** Percentage of DEGs per testis cell type between the *D. melanogaster* strains A4 and ISO1, shown relative to the *D. simulans* strain w^501^. DEGs are categorized based on whether they show consistent differential expression in both interspecific comparisons (A4 vs. w^501^ and ISO1 vs. w^501^) or are unique to only one of the comparisons. **e-h** Equivalent plots for the ovary. Genes were required to be expressed in at least 1% of cells with a minimum average expression of 0.01. Testis cell types: EC, epithelial cells; HC, hub cells; CC, cyst cells; ES, germline stem cells and early spermatogonia; LS, late spermatogonia; ESp, early spermatocytes; MSp, mid spermatocytes; LSp, late spermatocytes; MPSp, maturing primary spermatocytes; and S, spermatids. Ovary cell types: GSC/G1-2a, germline stem cells and germarium region 1 and 2a cells; G2a-2b, germarium region 2a and 2b cells; G2b-3, germarium region 2b and 3 cells; SPC, stalk and polar cells; FSC/preFCs, follicle stem cells and pre-follicle cells; EFC, early follicle cells; MFC, mitotic follicle cells stage 1-5; PMFC, post-mitotic follicle cells stage 6; V7, vitellogenic main-body follicle cells (MBFCs) stage 7; V8, vitellogenic MBFCs stage 8; V9-10A, vitellogenic MBFCs stage 9-10A; C12, choriogenic MBFCs stage 12; C14, choriogenic MBFCs stage 14; TCLC, terminal corpus luteum cells; SC, stretch cells; OSM, ovarian sheath muscle; and O, oviduct.

The two interspecific contrasts revealed overlapping but non-identical DEG sets (Fig. 3b,f), with 52.7% (1227/2328) and 46.4% (654/1408) shared between contrasts in the testis and ovary, respectively. For both tissues, and akin to our previous correlation analysis, the number of DEGs varied across cell types, peaking in the late and maturing primary spermatocytes in the testis, and in the germarium stages 1-2a and 2a-2b in the ovary (Fig. 3c,g; Supplementary Fig. 6). This sharply contrasts with the mitotic and somatic cell types in the testis and most somatic cell types in the ovary (*e.g.* the mitotic follicle and oviduct cell populations), which showed the lowest DEG counts between species. The number of DEGs per cell type was highly correlated across pairwise contrasts between strains (testis, minimum *r*^2^=0.78; ovary, minimum *r*^2^=0.81; Supplementary Table 8).

Expression divergence was also highly consistent at the cell type level. In the testis, 82.1% (1,007/1,227) of the DEGs were found in the same cell type, with an average consistency of 63.7% across cell types (minimum = 25.0%, in hub cells; maximum = 78.2%, in late spermatogonia) (Fig. 3d). The ovary showed a significantly higher consistency (92.5%, 605/654; 2STEP, χ^2^=37.07, d.f.=1, *P=*1.14×10^-9^), with an average of 74.5% (minimum = 62.5%, in stretch cells; maximum = 88.4%, in germarium 2a-2b) (Fig. 3h). Across both tissues, about 60% of consistent DEGs–genes differentially expressed in the same cell type for both interspecific contrasts–were differentially expressed in two or more cell types (Supplementary Fig. 7). Further, we detected a slight but significant tendency toward upregulation in *D. melanogaster* vs *D. simulans* for DEGs found only in testis versus those only in ovary (two-tailed Fisher’s exact test or FET, *P*=0.033, odds ratio=0.77). However, within each tissue, DEGs specific to one or shared between both tissues did not differ significantly in directional bias (FET, *P>*0.05) (Supplementary Fig. 8).

At the intraspecific level, we found substantially fewer DEGs (Fig. 3a,e), many of which also displayed interspecific differences. This overlap was significantly greater in the ovary (87.4% or 297/340) than in the testis (71.2% or 659/925; 2STEP, χ^2^=34.08, d.f.=1, *P=*5.28×10^-9^). After excluding overlapping DEGs, the global ratio of inter to intraspecific expression differences was also significantly higher in the ovary than in testis (14.07 vs 3.79; 2STEP, χ^2^=63.64, d.f.=1, *P=*1.5×10^-15^), denoting more relaxed functional constraints in the testis. This ratio varied substantially across cell types (Supplementary Fig. 9). By explicitly incorporating intraspecific variation, our experimental design avoids conflating standing expression variation with interspecific divergence, yielding a more accurate assessment of the selective constraints operating on transcriptome programs.

The increased resolution of snRNA-seq also uncovered fine-scale expression dynamics. For example, the gene *bol*, which encodes an RNA-binding protein essential for meiosis and spermatid differentiation ^46,47^, showed stable expression across strains and cell types (Fig. 4a), consistent with the action of stabilizing selection. Conversely, the gene *eIF4G1*, which encodes a translation initiation factor involved in spermatogenesis ^48^, was downregulated in both *D. melanogaster* strains relative to *D. simulans* (Fig. 4b), but only in late and maturing primary spermatocytes, and spermatids. This pattern is compatible with the action of lineage-specific selection, although additional intraspecific data from *D. simulans* and outgroup species are needed to substantiate this possibility. Equivalent patterns are found for key genes during oogenesis such as *orb* and *ovo* ^49,50^ (Supplementary Fig. 10a,b). Some genes showed more complex patterns of gene expression evolution, diverging in one or both *D. melanogaster* strains relative to *D. simulans*, with or without intraspecific changes (Fig. 4c).

**Figure 4.**
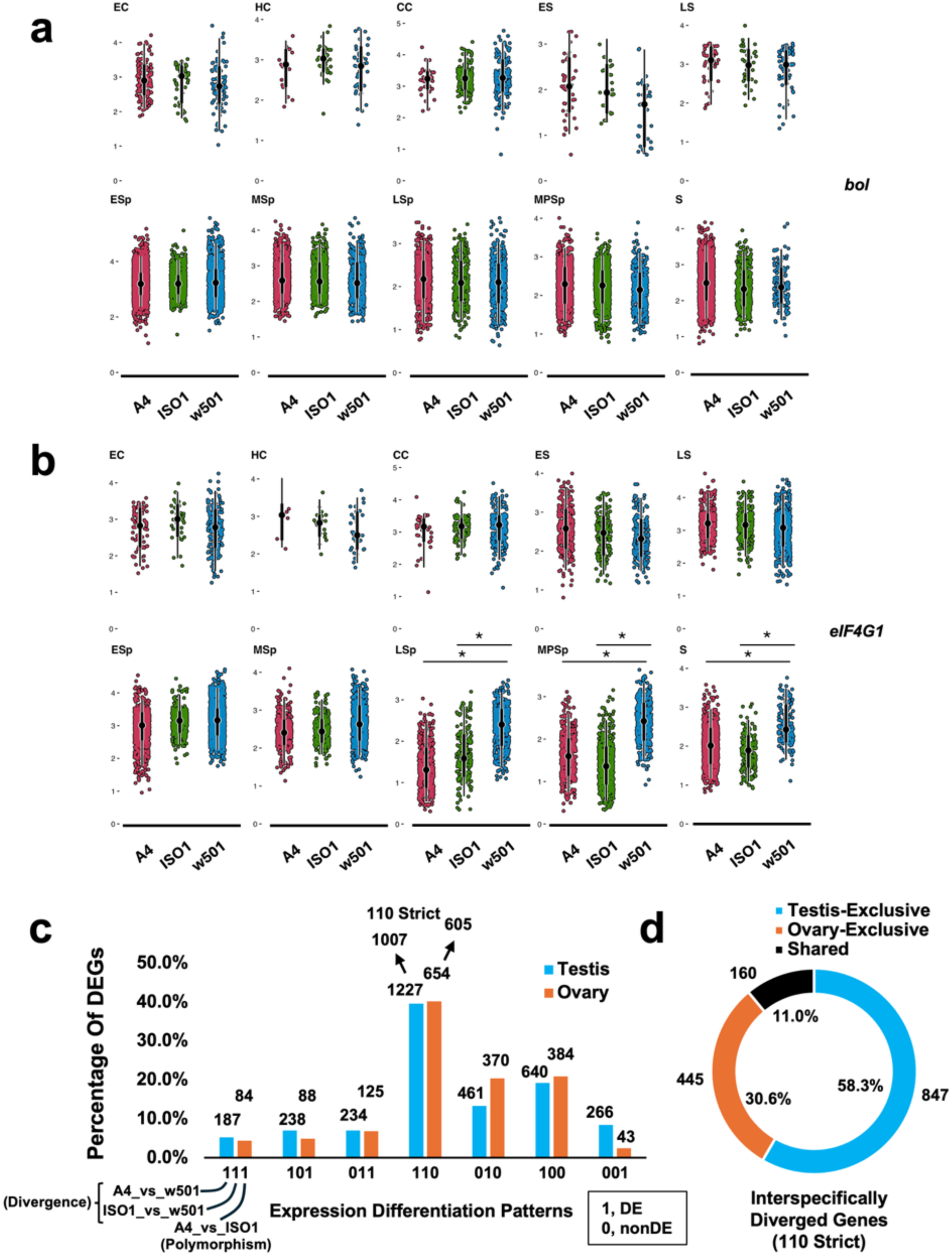
Expression differentiation at the cell type level in testis. **a** The gene *bol,* a non-DEG across cell types and strains, illustrates the absence of statistically significant differences in mRNA abundance among strains, a pattern consistent with functional constraints. **b** The gene *eIF4G1* exemplifies a gene that has diverged consistently at the interspecific level, but not at the intraspecific level. A4 and ISO1, *D. melanogaster* strains; w^501^, *D. simulans* strain. **c** Histogram showing the percentage and the number (above bars) of DEGs in at least one cell type in at least one of the pairwise contrasts among the three strains. The combination of presence (1) or absence (0) of expression differences in each of the three pairwise contrasts results in seven possible patterns of expression differentiation. The eighth pattern, corresponding to no significant expression differences in any comparison (000), is not shown. ‘110 Strict’ refers to the subset of DEGs entailing expression changes in at least one same cell type in both interspecific contrasts; most DEGs following a 110 pattern are part of this more stringent subset. **d** Donut chart showing the number and percentage of consistent DEGs at the interspecific level, exclusive to either testis or ovary, or shared between both tissues. Genes expressed in at least 1% of cells with a minimum average expression of 0.01 were considered. Testis cell types: EC, epithelial cells; HC, hub cells; CC, cyst cells; ES, germline stem cells and early spermatogonia; LS, late spermatogonia; ESp, early spermatocytes; MSp, mid spermatocytes; LSp, late spermatocytes; MPSp, maturing primary spermatocytes; and S, spermatids.

In total, we documented 1,452 DEGs consistently diverged across both interspecific contrasts in at least one common cell type in the testis and ovary, with only 11.0% shared across both tissues (Fig. 4d). Collectively, our findings suggest that the testis transcriptional program is more evolutionarily labile than that of the ovary, while providing enhanced resolution into how intra- and interspecific expression differences manifest at the level of individual cell types.

### Coexpression network modules diverge across species in a cell-type-specific manner

To determine whether gene expression divergence within individual cell types extended to broader coordinated changes at the network-level, we performed a coexpression network analysis (hdWGCNA ^51^). Our analysis revealed cell-type-specific modules of coexpressed genes encompassing 3,700 genes in the testis, and 4,730 genes in the ovary (Supplementary Table 9, Supplementary Data 5, 6). Roughly half of the constituent genes (1,642 in the testis and 2,655 in the ovary) were assigned to modules in more than half of the cell types, including known gametogenesis genes such as *polo* ^52^, *CycE* ^53^, and *mei-P26* ^54^. Conversely, a smaller set of genes (396 genes in the testis; 320 in the ovary) were uniquely assigned to modules in 20% or fewer cell types, highlighting potential cell type-specific roles (Supplementary Fig. 11).

To assess the functionality of these coexpression networks, we identified hub genes, defined as the top 10 genes within each module ranked by their intramodular connectivity, kME ^55^. These included known regulators of spermatogenesis and oogenesis, including *dj*, which is involved in mitochondrial differentiation within the flagellum during sperm individualization ^56^, *S-Lap8*, a protease-encoding gene crucial for spermatogenesis ^57^, and the previously mentioned *orb* and *ovo*, which are required for egg chamber formation and polarity establishment, and the activation of multiple maternally expressed genes during oocyte development, respectively ^49,50^. As expected, hub genes were enriched for GO terms tied to key aspects of germline development and gamete function, including spermatid development (GO:0007286), sperm motility (GO:0097722), and cytoplasmic translation (GO:0002181) (Supplementary Table 10).

Next, we examined whether the coexpression modules showed species-specific expression patterns by conducting a differential module eigengene (DME) analysis ^51^. We found that about half of the coexpression modules exhibited interspecific divergence across testis and ovary cell types, with at least one divergent module per cell type (Supplementary Table 9). In the testis, divergent modules were enriched during the later stages of spermatogenesis, consistent with our findings that species-specific transcriptional differences are most pronounced in the terminally differentiating meiotic cell types. A similar but non-significant trend was observed in the ovary, where terminal cell types—defined as the main body follicle cells—were compared against all other cell types (terminal cell types vs the rest, 2STEP; testis, χ^2^=4.62, d.f.=1, *P=*3.2×10^-2^; ovary, χ^2^=1.92, d.f.=1, *P=*0.17).

We further assessed the association between DEGs and the number of divergent coexpression modules across cell types, and found a positive correlation in the testis but not in the ovary (Spearman’s correlation coefficient; testis, ρ=0.782, *P=*7.5×10^-3^; ovary, ρ=-0.084, *P=*0.766). Notably, in the testis, 13 DEGs within interspecifically divergent modules are also found in an ancient spermatocyte protein interaction network conserved from *Drosophila* to humans ^58^. None of the 13 genes were part of our list of top 1% hub genes, and only the knock-down of one of them resulted in impairment of male fertility ^58^. These properties suggest that even genes part of this metazoan male germ cell network can accommodate transcriptional divergence at short phylogenetic distances with no detrimental consequences.

The DME and DEG results reinforced one another, providing a highly coherent picture of cell-type-specific expression divergence between *D. melanogaster* and *D. simulans*. For example, in late spermatocytes, the DME analysis revealed several divergent coexpression modules, particularly M1 and M3, with the former overexpressed in *D. melanogaster* and the latter in *D. simulans* (Fig. 5a). Examination of DEG distributions within these modules showed that in module M3, 29 constituent genes were upregulated in *D. simulans* (Fig. 5b, nodes in red), while none were downregulated, aligning with the overall pattern of network-level divergence. GO enrichment analysis of this module highlighted processes such as mitochondrial transport (GO:0006839), male gamete generation (GO:0048232), and spermatogenesis (GO:0007283) (Fig. 5d; Supplementary Table 11). In contrast, module M1 contained 27 genes upregulated in *D. melanogaster* and none downregulated (Fig. 5c, nodes in green). This module was enriched for processes such as microtubule cytoskeleton organization (GO:0000226) and nuclear division (GO:0000280) (Fig. 5e; Supplementary Table 11). Representative genes such as the microtubule-associated gene *jupiter* ^59^ and the sex differentiation gene *janB* ^60^ further illustrate the cell type-specific nature of these species-specific expression shifts (Fig. 5f,g). These results suggest that coexpression modules acquire species-specific regulatory programs during late spermatogenesis, reflecting functional specialization between *D. melanogaster* and *D. simulans*.

**Figure 5.**
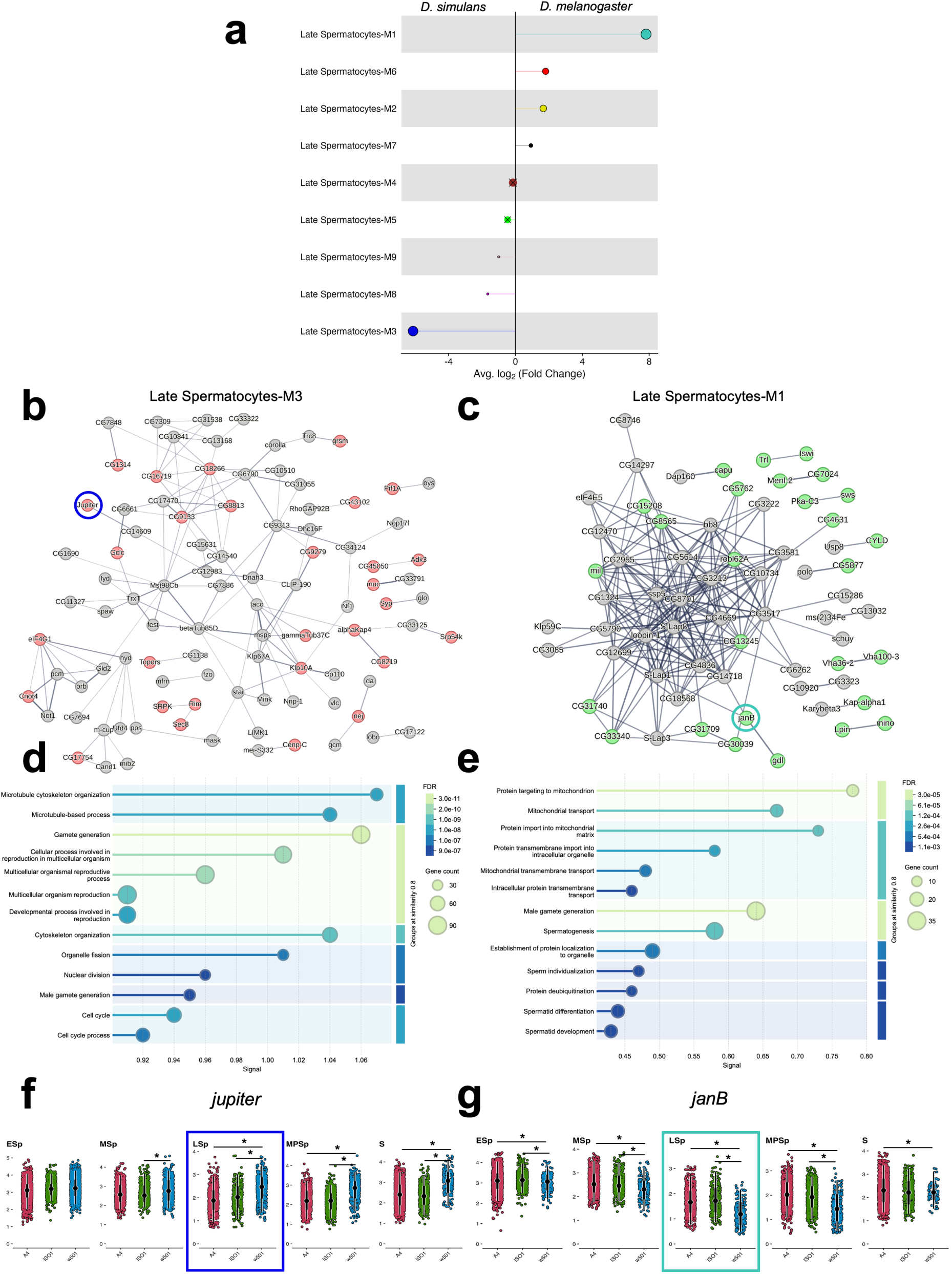
Coexpression module differentiation between *D. melanogaster* and *D. simulans* in late spermatocytes of the testis. **a** Differential module eigengene (DME) analysis showing the average log_2_ fold change in coexpression module activity between *D. melanogaster* and *D. simulans* in late spermatocytes. Modules M1 and M3 exhibit significant interspecific differences (minimum module size = 100 genes). **b, c** Coexpression network structure of the top 160 genes in Module M3 and Module M1 in late spermatocytes (medium confidence, *i.e.* 0.4, and 5% FDR). In Module M3 (**b**), red nodes denote genes downregulated in *D. melanogaster* relative to *D. simulans*, while in Module M1 (**c**), green nodes represent genes upregulated in *D. melanogaster* relative to *D. simulans*. Network edges reflect the level of confidence, and disconnected nodes are omitted. **d, e** Gene Ontology enrichment results for the Biological Process rubric for the top 500 genes in Module M3 and M1, respectively. **f, g** Expression patterns of the representative genes *jupiter* and *janB*, which are part of Module 3 (navy node in **b**) and Module 1 (turquoise node in **c**), respectively, across meiotic and post-meiotic testis cell types. The cell type of interest, late spermatocytes, is highlighted. A4 and ISO1, *D. melanogaster* strains; w^501^, *D. simulans* strain. Testis cell types: ESp, early spermatocytes; MSp, mid spermatocytes; LSp, late spermatocytes; MPSp, maturing primary spermatocytes; and S, spermatids.

Another example of cell-type specific network-level divergence was observed in stage 2a-2b germline cells of the ovary. The DME analysis uncovered key coexpression modules underlying early oogenesis divergence, with two modules, M1 and M2, displaying distinct coexpression patterns between species (Supplementary Fig. 12a,c). Module M1 was upregulated in *D. simulans* and was enriched for pathways related to germ cell development (GO:0007281) and oogenesis (GO:0048477) (Supplementary Fig. 13d; Supplementary Table 12). A notable example is *rhi* (Supplementary Fig. 12f), which encodes a germline-restricted heterochromatin protein essential for fertility and transposable element (TE) silencing ^61^, and has been implicated in female hybrid sterility between *D. simulans* and *D. melanogaster* ^62,63^. Conversely, Module M2 was overexpressed in *D. melanogaster* and predominantly enriched for cell cycle (GO:0007049) and related GO terms. One of the hub genes in this module, *mei-W68*, mediates double-strand break formation during meiosis ^64^ (Supplementary Fig. 12e,g; Supplementary Table 12). Notably, DEGs within both divergent modules exhibited typical connectivity relative to non-DEGs, indicating that divergence is not confined to peripheral genes, particularly for M1 where DEGs were in fact significantly more connected than non-DEGs (Kruskal-Wallis rank sum test; M1: χ²=17.94, d.f.=1, *P*=2.28×10^-5^; M2: χ²=1.11, d.f.=1, *P*=0.291). These findings demonstrate species-specific divergence in germline differentiation and cell cycle regulation during oogenesis, supported by both DME and DEG analyses that together reveal cell-type specific transcriptional changes between *D. melanogaster* and *D. simulans*.

### Contrasting patterns of functional constraints across cell types explain germline expression divergence

The differences in expression divergence across cell types—particularly the greater evolutionary lability of meiotic cells in the testis—suggest that functional constraints vary systematically across gametogenesis. To test this hypothesis, we examined whether genes expressed in different cell types differ in their pleiotropic signatures, given the established link between pleiotropy and the magnitude of functional constraints operating on a gene ^65–67^. Reduced expression breadth, a proxy for weaker functional constraint ^65,66^, was quantified using the tau index ^68^, calculated from FlyAtlas2 expression data ^69^ across gonadal tissues and cell types. In the testis, tau values of expressed genes differed significantly across cell types (Kruskal-Wallis test, *P*<1×10^-100^ for all three strains), increasing as spermatogenesis progressed, as confirmed by post-hoc tests (Fig. 6a and Supplementary Table 13). In the ovary, tau values also varied across cell types, with germline stem cells showing intermediate values and three somatic cell types—stalk and polar cells, and choriogenic MBFCs stage 12 and stage 14 cells (the latter two involved in the late stages of ovary differentiation)—showing the highest values (Fig. 6b and Supplementary Table 13). Overall, the meiotic cell types of the testis exhibited the highest tau values across all cell types and tissues, consistent with the relaxation of functional constraints, which is hypothesized to be associated with a more permissive chromatin environment found in these cell types ^70–72^.

**Figure 6.**
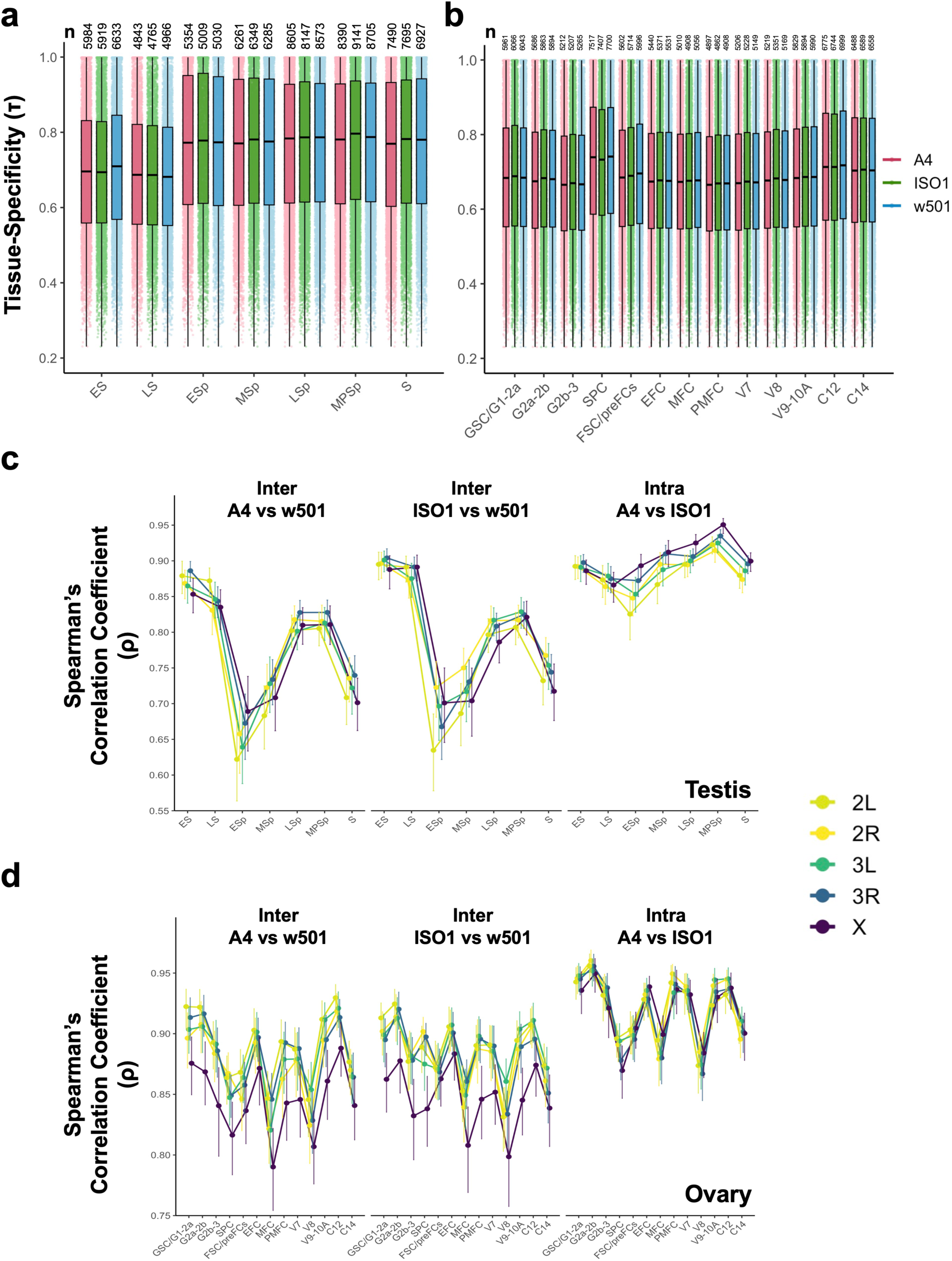
Expression breadth and chromosome expression differences across testis and ovary cell types. **a, b** Tau index (τ) distributions, a measure of expression specificity, across testis and ovary, respectively, for all expressed genes in specific cell types. Higher τ values indicate more restricted expression across cell types. Numbers above each boxplot denote the number of expressed genes per cell type. Only cell types directly involved in gametogenesis and with at least 100 cells per strain and 350 across all strains were included. Genes expressed in at least 1% of each cell type and with a minimum average expression of 0.01 are considered. Boxes represent the interquartile range around the median (black horizontal line) and whiskers extend to 1.5 times the IQR. Each point represents the correlation value of a particular gene, and values are grouped by strain and cell type. A4 and ISO1, *D. melanogaster* strains; w^501^, *D. simulans* strain. **c, d** Spearman’s correlation coefficient (ρ) of gene expression between strains across cell types in the testis and ovary, respectively, calculated separately for each major chromosome arm (*2*L, *2*R, *3*L, *3*R, and *X*). Error bars represent the 95% confidence interval across replicates. Correlation values above the 95th percentile within each cell type were omitted to reduce the impact of outliers in the expression correlation analyses.

Further, as expression breadth is correlated with phylogenetic gene age ^73^, we hypothesized that young genes (age categories B-E, *i.e.* genes that arose during the radiation of the genus *Drosophila*) would be enriched in late spermatogenesis stages, while older genes (*i.e.* age category A, predating the radiation of the genus *Drosophila*) would be overrepresented in mitotic cell types. Using a precisely curated gene age categorization in flies ^74^, we found broad support for these hypotheses (Supplementary Fig. 13). In contrast, the ovary revealed a very different picture with only the germline cell type G2b-3 showing enrichment for ancient genes, while the three aforementioned somatic cell types with higher tau exhibited a significantly elevated presence of younger genes from multiple age categories (Supplementary Fig. 14).

The rate of protein evolution serves as an additional proxy for assessing the magnitude of functional constraints. This rate covaries with expression divergence, especially for sex-biased genes ^16^, which are largely expressed in germline tissues ^12^. We used divergence and population polymorphism sequence data from *D. simulans*, and two different *D. melanogaster* populations, one from the ancestral range of this species and one from a subsequent colonization event to control for consistency, to calculate the rate of nonsynonymous to synonymous substitutions (ω)^75^. In the testis, ω of the expressed genes increased significantly in meiotic relative to mitotic cell types (Supplementary Fig. 15a and Supplementary Table 14). In the ovary, the choriogenic MBFCs at stage 12 exhibited a significantly higher ω than most other cell types (Supplementary Fig. 15d and Supplementary Table 15), and this pattern remained robust across all three strains. To disentangle the contribution of adaptive and nonadaptive mechanisms to differences in ω, we estimated ω_a_ and ω_na_, which measure the respective impact of these mechanisms ^76–78^. In the testis, the significant increase in the rate of protein evolution in meiotic cell types was primarily driven by significantly elevated ω_a_ values, reflecting a stronger effect of positive selection particularly in early meiotic cell types (Esp and MSp). In contrast, differences in ω between mitotic and late meiotic cell types (LSp and MPSp) were attributed mainly to non-adaptive evolutionary mechanisms (*i.e.* significantly higher ω_na_ values) (Supplementary Fig. 15b,c and Supplementary Table 14). In the ovary, however, we observed little to no variation in ω_a_ and ω_na_ across cell types (Supplementary Fig. 15e,f and Supplementary Table 15).

Collectively, these findings reveal markedly different patterns of functional constraint operating across the germline differentiation programs. Genes expressed during spermatogenesis show substantial variation in rates of protein evolution between mitotic, early meiotic, and late meiotic stages, shaped by differing contributions of adaptive and non-adaptive mechanisms. Moreover, compared to mitotic cells, meiotic cells in the testis tend to express genes that are more narrowly expressed and evolutionarily younger. In contrast, the ovary shows more uniform pleiotropic signatures across cell types.

### Genomic distribution of interspecifically divergent genes is biased in the female germline

The differences in functional constraints across cell types raise the question of whether a biased genomic architecture also underlies expression divergence. The hemizygosity of the *X* chromosome has been hypothesized to result in greater divergence at the coding and expression levels compared to autosomes ^79^. Although the evidence is mixed ^13,19,80–82^, fast-*X* effects have been primarily reported for male-biased genes in expression ^19,81^, many of which are preferentially expressed in the testis. We investigated how consistently divergent genes between species at the cell type level were distributed across the genome (Supplementary Fig. 16a). Consistent DEGs between species exclusive to the ovary, or shared across both tissues, deviated from the random expectation (chi-square goodness-of-fit test, χ^2^=51.35, d.f.=4, *P*_adj_*=*1.88×10^-10^, and χ^2^=26.74, d.f.=4, *P*_adj_*=*2.24×10^-5^, respectively). Post-hoc tests revealed that both categories of divergent genes were enriched on the *X* chromosome, with ovary-exclusive DEGs also significantly depleted from several autosomes (Supplementary Fig. 16b; Supplementary Table 16). In stark contrast, testis-exclusive DEGs showed a distribution not significantly different from the random expectation (chi-square goodness-of-fit test, χ^2^=5.96, d.f.=4, *P*_adj_*=*0.202) (Supplementary Fig. 16b). Additionally, to test the fast-*X* effect more broadly by including both DEGs and non-DEGs, we analyzed the correlation between expression levels per chromosome and cell type ^19,80,82^. In interspecific comparisons, we again found evidence that expression levels of *X-*linked genes were more divergent (*i.e.* lower Spearman’s correlation coefficients) than those of autosomal genes—but only in the ovary, in line with the above results (Fig. 6c,d; Supplementary Table 17). Unsurprisingly, the intraspecific test showed no differences across chromosomes, and all patterns were recapitulated at the pseudo-bulk level (Supplementary Fig. 17a).

Both adaptive and non-adaptive mechanisms can generate a fast-*X* effect ^83^. To distinguish between these possibilities, we assessed the strength of purifying selection on gene expression by analyzing the correlation of gene expression between the testis and ovary in each strain across chromosomes at the pseudo-bulk level ^19^. Across strains and species, the *X* chromosome consistently exhibited higher testis-ovary correlation values than most autosomes (Supplementary Fig. 17b). These findings suggest that the pronounced expression divergence on the *X* is unlikely to result from relaxed functional constraints, instead pointing to positive selection as a more plausible driver of the observed fast-*X* effect.

Overall, our findings challenge traditional views from previous *Drosophila* studies ^19,81,82^. The enrichment of DEGs exclusive to the ovary and those shared between tissues on the *X* chromosome points to a substantial, persistent role for positive selection acting on *X-*linked genes involved in the reproductive fitness of females—the sex in which the *X* chromosome resides two-thirds of the time in any given generation.

## DISCUSSION

Our analysis of 106,328 testis and ovary nuclei from three *Drosophila* strains—to our knowledge, the largest snRNA-seq dataset for *Drosophila* gonads—provides novel insights into cell type-specific transcriptional divergence at short phylogenetic distances. Traditional bulk transcriptomic studies in *Drosophila* have lacked cell type resolution ^13,16,17,20,84,85^, limiting inferences about fine-scale expression differences. We identified numerous key genes involved in spermatogenesis (*Jupiter*, *janB,* and *eIF4G1* ^59,60,86^) and oogenesis (*ovo*, *rhi*, and *mei-W68* ^49,61,64^) that display interspecific expression differences in a cell-type-specific manner, extending similar observations from mammalian gonads ^9^ and other tissues ^10,41,87–90^. Our results also reveal new insights into gene functionality. For example, *ovo*, known for its central role in oocyte development ^49^, shows species-specific expression in late-stage follicle cells, suggesting a somatic role consistent with isoform-specific activity ^91^.

Transcriptome evolution during spermatogenesis is especially dynamic, with divergence concentrated in meiotic and post-meiotic germ cells. These stages harbor more DEGs and divergent coexpression networks than earlier stages, suggesting reduced constraints and increased opportunity for regulatory divergence. Our findings align with prior studies on sectioned testis from *D. melanogaster* and *D. simulans* ^92^ and mammalian snRNA-seq that identify late spermatogenesis as a hotspot of expression divergence ^9^. In mammals, this has been linked to narrower spatiotemporal expression profiles and a greater prevalence of evolutionarily young genes ^9^, facilitated in part by a more permissive chromatin state ^70–72^. These factors may allow both neutral and adaptive expression changes to accumulate—neutral changes due to weak or no purifying selection ^93^, and adaptive changes in response to different selective pressures, most notably sperm competition ^72,94^.

We also found evidence of relaxed constraint as spermatogenesis progresses in *Drosophila*. Evolutionarily younger genes with narrower expression breadth and faster rates of sequence evolution are more prevalently expressed at later stages, which also display lower ratios of interspecific to intraspecific expression changes. Consistent with these patterns, our network-level analyses identified mitochondrial transport and microtubule-based processes—both essential for germ cell development ^95,96^—as major targets of transcriptome divergence in late spermatogenesis. This divergence may reflect species-specific adaptations shaped by sexual selection, particularly sperm competition ^97^. Notably, sperm length—a key factor in sperm competition—depends on the microtubule cytoskeleton for elongation ^96^, a trait that differs markedly between *D. melanogaster* and *D. simulans* ^98^. Together, these findings support a model of accelerated transcriptome evolution in late spermatogenesis, driven by a combination of adaptive and non-adaptive evolutionary mechanisms.

In the ovary, although overall divergence is more limited than in the testis (605 DEGs in the ovary vs 1,007 DEGs in the testis), cell-type-specific differences are still prominent. Genes expressed in the ovary tend to show more uniform evolutionary properties across cell types—older age, broader expression, and slower sequence evolution—reflecting stronger pleiotropic properties and therefore under stronger evolutionary constraints. However, the global ratio of interspecific to intraspecific expression changes is significantly higher in the ovary than in the testis, suggesting a stronger role for adaptive mechanisms driving the interspecific transcriptome divergence of the ovary. Among the most divergent cell types are terminally differentiating main-body follicle cells, which provide structural and regulatory support to the oocyte ^99^, and early-stage germline cells. For example, in stage 2a-2b germline cells, which contain a mix of meiotic and nurse cells that have exited meiosis ^100^, we identified divergent coexpression modules associated with TE suppression, cell cycle processes, and meiotic progression. This mirrors observations in mammals, where species-specific divergence in early oogenesis is linked to variation in cell cycle timing ^101^. The presence of meiotic gene expression in early germline cells of the germarium may help explain why divergence is detectable at these early stages ^102^. These observations suggest that fine-scale regulation of mitotic-to-meiotic transitions—or meiosis itself—may be an important target of expression divergence in the ovary, as in the testis, with potential implications for oocyte quality and reproductive success. Together, our findings challenge the view of the ovary as transcriptionally static during evolution and reveal that even core biological processes in the ovary can undergo cell-type-specific divergence.

By integrating gene- and network-level analyses, our study represents a pioneering effort in comparative transcriptomics of insects, revealing a more nuanced view of expression divergence during spermatogenesis and oogenesis at single-cell resolution. Our results support a more granular model of transcriptome evolution than previously recognized—one that reflects distinct biological demands and exposure to selection mechanisms ^103,104^. Lastly, we provide a valuable online resource for further exploring interspecific expression divergence during gametogenesis.

## ACKNOWLEDGEMENTS

We thank Brandon Gaut, Anthony Long, Ana Llopart, and Kyoichi Sawamura for comments on early versions of this manuscript. We also thank the Genomic High Throughput Facility and the Research Cyberinfrastructure Center at the University of California Irvine for facilitating our sequencing experiments and computational analyses, respectively.

## AUTHOR CONTRIBUTIONS

I.E.H. and J.M.R. conceived and designed the study, performed downstream analysis and interpreted the results. I.E.H. performed most bioinformatic analyses and supervised and contributed to dissections. S.D. and I.E.H. performed nuclei isolation, library preparation, and sequencing. E.M.L., C.G.-C., and T.S. carried out dissections and tissue collection. J.C. and I.E.H. built a shiny app for visualizing the snRNA-seq data. V.S. provided input on study design and data interpretation. J.M.R. supervised the project and contributed specific analyses. I.E.H. and J.M.R. co-wrote the manuscript. All authors reviewed and approved the final manuscript.

## METHODS

### Fly husbandry

Two strains of *D. melanogaster* (A4 and the reference strain ISO1), and the strain w^501^ of *D. simulans* were used. Flies were reared on a standard dextrose-cornmeal-yeast medium at room temperature (∼25 °C) under continuous light. All manipulations of flies were performed under CO_2_ anesthesia.

### Gonadal tissue dissection

Testis dissections were performed on 1-2-day-old virgin males and ovary dissections on 3-day-old virgin females using fine-tipped forceps in cold Schneider’s Drosophila Medium (ThermoFisher, #21720024). The selected age prevented the overrepresentation of mature sperm that characterizes older males ^7^, and ensured that females were fully fertile and receptive^105^. Dissected tissues were immediately transferred to 200 μL of Schneider’s Medium in nuclease-free 1.5 mL Eppendorf tubes, pooled (10-40 tissues per tube), and kept on ice for less than 2 h. Separate petri dishes were used for dissections of different strains and tissues to avoid cross-contamination and ensure sample integrity. After dissections were completed, samples were flash-frozen by submerging the tubes in liquid nitrogen and immediately stored at -80 °C for long-term preservation. To maintain RNA integrity, samples were thawed only prior to cells isolation. Two biological replicates for each dissected tissue and strain were generated. In total, we dissected 1200 pairs of testes (200 pairs per biological replicate) and 240 pairs of ovaries (40 pairs per biological replicate).

### Nuclei isolation

Nuclei isolation from *Drosophila* ovary and testis was performed using a two-step dissociation process. Initially, tissues were incubated in HBSS Buffer containing 2.5 mg/mL collagenase (Invitrogen, #17018-029) at 37°C for 15 min. The tissues were then homogenized in EZ Lysis buffer (Sigma-Aldrich, #NUC101-1KT) and incubated on ice for 10 min, followed by filtration through a 70 μm filter. The filtered homogenate was centrifuged at 750 g for 5 min at 4°C and resuspended in 1mL of lysis buffer. After a second centrifugation, the nuclei were incubated in Nuclei Wash and Resuspension buffer (1xPBS, 1% BSA, 0.5U/μL RNase inhibitor) for 5 min. To remove debris, a debris removal solution (Miltenyi Biotec, #130-109-398) was added to the nuclei suspension, and the mixture was centrifuged at 3,000g for 10 min at 4°C. The purified nuclei were fixed and permeabilized using the Nuclei Fixation Kit (Parse Biosciences) and cryopreserved in DMSO until library preparation.

### Preparation of snRNA-seq library and sequencing

Libraries were prepared using the EVERCODE^TM^ WT V2 kit (Parse Biosciences) and quantified with the Qubit dsDNA HS assay kit (Invitrogen, #Q32851). The average fragment length of each sub-library was measured using the D5000 HS kit (Agilent, #5067-5592, #5067-5593). Finally, the eight sub-libraries were sequenced on the Illumina Novaseq 6000 S4 platform using paired-end sequencing at the UCI Genomics Research and Technology Hub, with an estimated sequencing depth of 50,000 read pairs/nuclei.

### snRNA-seq read alignment

Raw sequencing data was assessed using FastQC (version 0.11.9) ^106^ to evaluate read quality, sequence composition, and potential sequencing artifacts. Read alignment was performed against the respective genome assembly of each *Drosophila* strain: ISO1 ^107^, A4 ^32^ and w^501^ ^108^. To facilitate comparative genomic analysis, we employed Liftoff (version 1.6.3) ^109^ to annotate the A4 genome by mapping gene annotations from the ISO1 reference genome to the A4 assembly using default parameters. We filtered each annotation in the GTF file to retain protein-coding genes only using 10X Genomics’ CellRanger mkgtf (version 8.0.1) ^110^ prior to creating the references using the ‘mkref’ mode of Parse Biosciences’ split-pipe pipeline (version 1.1.2) ^111^. Reads from each of the eight sub-libraries were demultiplexed by strain and aligned independently to the three respective references using the ‘all’ mode of split-pipe. The alignments were configured to incorporate each sample’s designated wells, in addition to specifying the V2 chemistry of the library preparation protocol and a post_min_map_frac of 0.01. Lastly, we combined the resulting alignments from the eight sub-libraries for each sample using the ‘comb’ mode of split-pipe to generate the final count matrices.

### Quality control, normalization, and integration

We mapped orthologs from *D. melanogaster* to *D. simulans* using previously delineated calls ^45^ and replaced the *D. simulans* gene names while retaining the genes without any one-to-one mapping. We combined the data from our 2 biological replicates to create one Seurat object for each tissue and strain, applying a stringent quality control criteria with the following thresholds: min.features = 200, min.cells = 3, and a minimum nCount_RNA of 300 for testis and 500 for ovary (version 5.1.0) ^112^. We removed nuclei with mitochondrial reads above 5% and limited the dataset to nuclei with a maximum of 7,500 expressed genes and 100,000 counts to remove outliers. These thresholds were implemented while inspecting violin and feature scatter plots to assess the distribution of the metrics across samples. Each of these datasets was then log-normalized (NormalizeData) and scaled (ScaleData) before obtaining a tissue- and strain-specific UMAP using Seurat’s built-in functions (RunPCA, FindNeighbors, FindClusters, RunUMAP).

DoubletFinder (version 2.0.4) ^113^ was employed to minimize the potential effect of doublets by using the recommended iterative parameter sweeping approach to identify the optimal pK (principal component) value for each Seurat object. We assumed a default doublet formation rate of 3% (https://support.parsebiosciences.com/hc/en-us/articles/360053107311-What-is-the-expected-doublet-rate) and used a homotypic doublet proportion estimation based on initial clustering results. Nuclei classified as doublets were removed from downstream analyses.

The post-filtered Seurat objects were merged into a single object for each tissue, and normalization (NormalizeData) and variable feature identification (FindVariableFeatures) were performed for each layer independently. The cross-strain layers were integrated using Harmony (version 1.2.1) ^44^, which performed the best among other tools tested (LIGER ^114^, RPCA - https://satijalab.org/seurat/articles/integration_rpca.html, CCA ^115^, scVI ^116^), and is also recommended for closely related species, carrying out species mixing while preserving biological heterogeneity ^117^.

### Clustering and cell type annotation

The integrated layers were subsequently joined for each tissue, and an ElbowPlot was used to determine the number of PCs for the FindNeighbors command for both testis (dims = 1:15) and ovary (dims = 1:20). We used clustree (version 0.5.1) ^118^ for interrogating the clusters at increasing resolution before running FindClusters on the testis (resolution = 1.2) and ovary (resolution = 1) Seurat objects. We generated a combined UMAP for each tissue using the harmony reductions and visualized the plots using scpubR (version 2.0.2) ^119^. Cell type annotations were inferred using known marker genes specifically expressed in each cell type (Supplementary Table 4 and Supplementary Fig. 3,4), as documented in recently published single-cell and single-nucleus RNA-seq studies of the gonadal tissue of *Drosophila* (testis ^7,22–24^; ovary ^7,25,26,28^).

Several control analyses were implemented to confirm expected species clustering and validate our cell type annotation. First, we performed a pseudo-bulk analysis and generated an MDS plot for each cell type of each tissue, confirming that biological replicates clustered by strain. Second, the proportions of different cell types were compared across strains using pairwise Pearson’s correlations, ensuring that no unexpected or artifactual clustering patterns arose (Supplementary Note 1). Lastly, a trajectory analysis using Monocle3 (version 1.3.5) ^120^ was performed to demonstrate that the annotated cell types were arranged in the expected developmental sequence of gametogenesis for each tissue (Supplementary Fig. 1).

### Principal component analysis of cell-type pseudo-bulks

To examine expression differences within each tissue, we aggregated raw gene expression counts by cell type and sample to generate pseudo-bulk profiles. We combined these profiles into a single matrix with corresponding cell type and sample metadata, removed genes with zero counts across all pseudo-bulks, and applied variance-stabilizing transformation using the rlog function in DESeq2. We then performed principal component analysis on the transformed matrix and visualized the top two components to assess variation between cell types and strains (Fig. 2a-b).

### Correlation of gene expression

To assess the degree of gene expression correlation across *Drosophila* strains, we identified orthologous genes present in all three strains (ISO1, A4, and w^501^). We extracted the average gene expression levels of all 1-to-1 orthologous genes between *D. melanogaster* and *D. simulans* and generated a correlation matrix using Pearson correlation coefficients, which was visualized using a heatmap.

### Differential gene expression analysis

We performed pairwise differential gene expression analyses for each interspecific (A4 vs w^501^; ISO1 vs w^501^) and intraspecific (ISO1 vs A4) comparison for testis and ovary using MAST (version 1.28.0) ^121^. Differential expression was evaluated for each gene and cell type by comparing expression levels between two given strains under the following criteria: |log2 fold change| ≥ 1 and FDR ≤ 1%. We further corrected for multiple testing across the three comparisons using the Benjamini-Hochberg method (FDR ≤ 1%) and classified genes as differentially expressed (DE) or non-differentially expressed (nDE). Genes that did not meet a minimum expression threshold of min.pct = 0.01 in a given cell type for both strains being compared were excluded from the analysis, thereby mitigating possible artifacts associated with low expression.

To categorize the differential expression patterns across the three pairwise comparisons, we adopted a systematic approach to characterize expression differentiation patterns in the testis and ovary using custom in-house Python scripts. We implemented an encoding system where each gene received a three-digit binary code representing its differential expression status across these comparisons (A4 vs w^501^; ISO1 vs w^501^; ISO1 vs A4) for each tissue. Each digit can either be ‘0’ for an nDE gene or ‘1’ for a DE gene in at least one of the cell types considered (10 for the testis and 17 for ovary, respectively). Thus, interspecifically diverged genes had a code of ‘110’, which was compared between testis and ovary to investigate tissue-specific gene expression evolution. A similar 10- or 17-digit trinary code (0, no difference; +1, overexpression in strain 1 vs strain 2; and -1, underexpression in strain 1 vs strain 2) was applied for summarizing patterns of differential expression across cell types within each comparison for the testis and ovary, respectively.

### Weighted gene co-expression network analysis (hdWGCNA)

To identify modules of significantly co-expressed genes, we performed weighted gene co-expression network analysis (WGCNA) using the hdWGCNA framework (version 0.3.01) ^51^. The analysis was carried out independently for each cell type, incorporating cells from all three strains, focusing exclusively on genes shared across strains and expressed in at least 5% of cells. Metacells were constructed by grouping cells with nearest-neighbor (k = 25) aggregation using the Harmony-reduced space. Metacells shared by more than 15 cells were excluded, groups with fewer than 50 cells were discarded, and the resulting metacell expression matrix was log normalized. The “signed” network type was used for optimizing the soft-thresholding power and maximize the scale-free topology fit. A topological overlap matrix (TOM) was computed for each cell type, and hierarchical clustering with dynamic tree cutting (deepSplit = 4, minimum module size = 25) was applied to identify co-expression modules. Module eigengenes (MEs) were calculated for each module and hub genes within each module were identified based on eigengene connectivity (kME) by ranking genes by kME. Modules assigned to the “grey” category, indicating unassigned genes, were excluded from downstream analyses.

To assess interspecies differences in module activity, we performed differential module eigengene (DME) analysis using hdWGCNA’s ‘FindDMEs’ function. Module eigengene expression profiles were compared between *D. melanogaster* and *D. simulans*, applying min.pct = 0.01 to exclude lowly expressed genes. We identified significantly divergent modules for each cell type using the Wilcoxon rank-sum test and compared the constituent genes of these divergent modules with the differentially expressed genes to examine congruency between network-level and gene-level patterns of expression divergence.

### Functional enrichment analysis

We analyzed gene ontology (GO) term enrichment among genes present in co-expressed gene modules for each cell type and tissue using clusterProfiler (version 4.0) ^122^ with genome-wide annotations from org.Dm.eg.db (version 3.18.0) ^123^. Enrichment results were restricted to the top 500 genes per module and we applied a 5% FDR ^124^. To identify shared and cell-type-specific functional enrichments, we used the compareCluster function in clusterProfiler to compare GO term enrichment across modules.

### Genomic features

We examined several functional and genomic features in the context of expression divergence between *D. melanogaster* and *D. simulans.* Chromosomal locations and sequence coordinates were retrieved from FlyBase ^125^. For expression breadth, we obtained gene expression values across 31 tissues and body parts from adult individuals of both sexes as in FlyAtlas2 ^69^, and calculated the tissue-specific index tau (τ) (Yanai et al. 2005). The tau index ranges from 0 to 1 with lower and higher values indicating broader and narrower expression, respectively.

We retrieved the phylogenetic age of genes inferred under a parsimonious framework based on syntenic alignments among 20 fly species, including the species *Scaptodrosphila lebanonensis* and *Bactrocera dorsalis*, which are outgroups to the genus *Drosophila* ^74^. The five age classes considered correspond to different tree branches in the original publication: class A corresponds to genes originated in branches -1 and -2, *i.e.* those present before the *Drosophila* radiation; class B corresponds to branch 0, *i.e*. those that arose prior to the split between the *Drosophila* and *Sophophora* subgenera; class C corresponds to branches 1 and 2, *i.e.* to those genes originated in the lineages to the *D. willistoni* and *D. pseudoobscura* species groups; class D corresponds to branch 3, *i.e.* to those genes originated during the radiation of the *D. melanogaster* species group; and class E corresponds to branches 4-6, *i.e.* to those originated during the evolution of the *D. melanogaster* species subgroup, which includes *D. melanogaster* and *D. simulans*.

Gene sequence evolution was analyzed through a modified version of the McDonald and Kreitman test, the extended McDonald and Kreitman test ^126^, which controls for low frequency polymorphisms and differentiates neutral from weakly deleterious variants, thus minimizing the problem of underestimating α, *i.e.* the proportion of substitutions fixed by positive selection (α) ^78^. Synonymous and nonsynonymous nucleotide substitutions were obtained from the comparison of sequences involving *D. simulans,* 197 lines from a *D. melanogaster* African population (Zambia), and 205 lines from a North American *D. melanogaster* population (Raleigh, North Carolina) using the iMKT R package ^127^. From the iMKT site, we also downloaded the allele frequency spectrum. To estimate the number of neutral segregating sites in the non-synonymous category, we applied a minimum 5% frequency threshold to remove slightly deleterious mutations^127^. Alpha was calculated considering the corrected estimates, which allowed us to calculate the rate of protein evolution (ω), the rate of adaptive evolution (ωa), and the rate of non-adaptive evolution (ωna) ^76,77^.

### Statistical Analyses

Two-sample test for equality of proportions with continuity correction, two-tailed Fisher’s exact test, chi-square goodness-of-fit, chi-square test of independence, analysis of residuals, Kruskal-Wallis rank sum, post-hoc Dunn’s pairwise tests, regression analysis, linear modelling, and Benjamini-Hochberg correction for multiple testing were conducted using built-in functions in R (version 4.3.2) ^128^.

